# Poly(acrylamide-co-N,N’-methylene bisacrylamide) monoliths for high peak capacity hydrophilic interaction chromatography high-resolution mass spectrometry of intact proteins at low trifluoroacetic acid content

**DOI:** 10.1101/2021.07.14.452317

**Authors:** Marta Passamonti, Chiem de Roos, Peter J. Schoenmakers, Andrea F.G. Gargano

**Affiliations:** Van’t Hoff Institute for Molecular Sciences, University of Amsterdam, Amsterdam, The Netherlands; Centre for Analytical Sciences Amsterdam, Science Park 904, 1098 XH Amsterdam, The Netherlands

**Author notes:** Corresponding Authors; Mailing Address: Marta Passamonti, Faculty of Science, Van ‘t Hoff Institute of Molecular Science (HIMS), P.O. Box 94157, 1090GD Amsterdam, The Netherlands.; Mailing Address: Analytical Chemistry, Center for Analytical Sciences Amsterdam, Science Park 904, Building C2, room 258, Universiteit van Amsterdam, 1098 XH Amsterdam, The Netherlands.

## Abstract

In this study, we optimized a polymerization mixture to synthesize polyacrylamide-co-N,N’-methylene bisacrylamide monolithic stationary phases for hydrophilic-interaction chromatography (HILIC) of intact proteins. Thermal polymerization was performed, and the effects of varying the amount of crosslinker and the porogen composition on the separation performance of the resulting columns were studied.

The homogeneity of the structure and the different porosities were examined through scanning electron microscopy. Further characterization of the monolithic structure revealed a permeable (*K_f_* between 2.5 × 10^−15^ and 1.40 × 10^−13^ m^2^) and polar stationary phase suitable for HILIC. The HILIC separation performance of the different columns was assessed using gradient separation of a sample containing four intact proteins, with the best performing stationary phase exhibiting a peak capacity of 51 in a gradient of 25 min.

Polyacrylamide-based materials were compared with a silica-based particulate amide phase (2.7 μm core-shell particles). The monolith has no residual silanol sites and, therefore, fewer sites for ion-exchange interactions with proteins. Thus, it required lower concentrations of ion-pair reagent in HILIC of intact proteins. When using 0.1% of trifluoroacetic acid (TFA) the peak capacities of the two columns were similar (31 and 36 for the monolithic and packed column, respectively). However, when decreasing the concentration of TFA to 0.005%, the monolithic column maintained its separation performance and selectivity (peak capacity 26), whereas the packed column showed greatly reduced performance (peak capacity 7), lower selectivity, and inability to elute all four reference proteins. Finally, using a mobile phase containing 0.1% formic acid and 0.005% TFA the HILIC separation on the monolithic column was successfully hyphenated with high-resolution mass spectrometry. Detection sensitivity for protein and glycoproteins was increased and the amount of adducts formed was decreased in comparison with separations performed at 0.1% TFA.

## INTRODUCTION

Monolithic stationary phases are single porous pieces of stationary phase obtained through thermal- or photo-polymerization of a mixture consisting of monomer, crosslinker, porogens and initiator. These stationary phases have been extensively investigated, because of their inherent high permeability, low resistance to mass transfer, fast and simple preparation in small formats^1–5^ and a plethora of possible chemistrIes and retention mechanisms^6–8^.

In the mid 1990s, the first acrylamide-based monoliths were developed. Hjertén and co-workers prepared acrylamide monolithic stationary phases based on a protocol that involved the dispersion of polar monomers and crosslinkers in aqueous solutions. They obtained what they called compressible beads for LC separations.^9–11^ Later, this protocol was optimized by the group of Freitag, who added dymethylformamide (DMF) as a porogen to harden the monolithic structure in capillaries used for electrochromatography applications.^12^ In 1997 Xie et al. studied an alternative approach to generate more rigid materials, co-polymerizing acrylamide and methylene bisacrylamide (MbA) in dimethyl sulfoxide (DMSO) and 2-heptanol as organic solvents.^13^ Materials with pore size up to 1000 nm were prepared by adjusting the synthesis conditions. No application of these materials for chromatographic purposes was reported. However, similar polymerization mixtures were later used for the production of immobilized enzyme reactors^14^ and capillary electrochromatography^15^ columns.

To the best of our knowledge the first applications of hydrophilic interaction chromatography (HILIC) to the study of intact proteins involved the separation of histone proteins on a weak-cation-exchange column^16–18^ and later the separation of membrane proteins.^19,20^ Zhang ^21^, Pedrali^22^, Lauber^23^ and Periat^24^ et al. discussed the unique selectivity of HILIC in the separation of glycoforms of sub-units and intact glycoproteins. These results have drawn attention to the use of HILIC-MS, with recent applications reporting high-resolution separations of neo-glycoproteins^25^, biopharmaceuticals^26–29^, biotechnological products^30^ and serum immunoglobins^31^. For these proteins, HILIC resolved proteoforms that would co-elute in RPLC.

The majority of the separations of intact proteins and subunits by HILIC use silica particles with amide selectors and mobile phases based on acetonitrile, water and acidic ion-pairs, such as trifluoroacetic acid (TFA). Negatively charged ion-pair reagents are thought to neutralize the polar basic groups present in proteins, improving peak shapes and increasing the contribution of neutral polar groups, such as glycans, to retention. However, TFA and other ion-pair reagents greatly reduce MS sensitivity and can lead to gas-phase ion adducts, which complicate the mass spectra.^32^

The residual acidity of silica based materials (free silanols) is thought to be a source of non-glycan-specific ion-exchange interactions and, therefore, increases the need for ion-pair reagents in the mobile phase to enhance the separation efficiency and reduce peak tailing for glycoforms. To reduce the amount of TFA used in HILIC separations of proteins Wirth and co-workers^21^, bonded a polyacrylamide layer on non-porous silica particles of 700 nm size packed in a 2.1-μm (internal diameter) ID column. With this material they obtained baseline separation of the five proteoforms of ribonuclease B, lowering the amount of TFA to 0.05% and adding 0.5% formic acid (FA). However, only the use of polymeric materials that are not based on silica, but, for example, exclusively on polyacrylamide, can fully exclude ion-exchange interaction with silanols. To date, no fully organic material for HILIC of intact proteins has been reported.

In this study, we set out to synthesize and optimize an acrylamide-based monolithic stationary phase for protein separations in HILIC. We aimed to explore the effects of varying the reaction mixture (monomer, crosslinker and porogens) and to characterize the different capillary columns in terms of permeability, hydrophilicity, morphology and porosity. Our aim was to obtain efficient columns for HILIC separations of intact proteins and proteoforms, which could be operated with very low concentrations of TFA as ion-pair reagent, so as to greatly enhance the sensitivity of MS detection in comparison with capillary columns packed with commercial silica-based amide stationary phases.

## EXPERIMENTAL SECTION

### MATERIALS

Acrylamide (AA, electrophoresis grade, 99%), *N,N’*-methylenebisacrylamide (MbA, 99%), 3-(trimethoxysilyl)propyl methacrylate (γ-MAPS, 98%), 2,2’-azobisisobutyronitrile (AIBN, 98%), 1-octanol (OctOH, 99%), dimethyl sulfoxide (DMSO >99.9%), toluene, sodium hydroxide (NaOH), lysozyme from chicken egg white (Lys), carbonic anhydrase from bovine heart (CA, >90%), myoglobine from equine heart (Myo, >90%), cytochrome C (CC), bovine serum albumin (BSA), ribonuclease A from bovine pancreas (RnA), ribonuclease B from bovine pancreas (RnB, ≥80%), transferrin from human serum, (Tf), cytosine (Cyt), trifluoroacetic acid (TFA, ≥99%), and formic acid (FA, analytical grade >98%) were purchased from Sigma Aldrich (St. Louis, Missouri, United States). Methanol (MeOH), acetonitrile (ACN) were purchased from Biosolve (Valkenswaard, The Netherlands). Hydrochloric acid 37% (HCl) was obtained from Acros (Geel, Belgium). High-purity (HP) water (18.2 MΩcm) was produced by a Sartorius (Göttingen, Germany) Arium 611UV Ultrapure-Water System. The capillary (0.20 mm ID, 0.36 mm OD) was purchased from CMScientific (Silsden, UK). Glass-lined tubing (300 mm length × 0.80 mm ID) was purchased from VICI (Houston, TX)

### PREPARATION OF POLY(ACRYLAMIDE-CO-N,N’-METHYLENEBISACRYLAMIDE) MONOLITHIC COLUMNS

Monolithic capillary columns were optimized for separations of intact proteins. To covalently attach the monolith to the inner wall surface, the wall of the fused-silica capillary had to be modified prior to the polymerization reaction. The surface was etched using NaOH and HCl and subsequently silanized using a 20% (v/v) γ-MAPS solution in toluene, as described by Courtois *et al*.^33^ Thereafter, the capillary was flushed with toluene and dried with nitrogen. Several acrylamide-based monoliths were synthesized sinside the capillaries tarting from polymerization conditions similar to those described by Xie *et al.*^13^, using the recipes summarized in Table 1. The most-difficult step was to completely dissolve the monomer and the crosslinker in the porogens. In order to solubilize the monomers, the MbA was added to the porogens and sonicated for 1 h at a temperature between 30 and 40°C. Once the crosslinker was dissolved, we added the monomer and kept sonicating for 45 min at the same temperature. Finally, the AIBN (1% w/w with respect to the monomers) was added and the mixture was vortexed. Thereafter, the silanized capillaries were filled with the polymerization mixture, and the ends were closed using rubber pieces. Polymerization took place for 24 h at 60°C in a thermostated glass tube. Finally, the monolithic capillary columns were thoroughly flushed with MeOH.

**Table 1.** Composition of the polymerization mixtures (monomers and porogens) used to prepare the different acrylamide-based columns (the abbreviations of monomers and porogens are isted in the Materials section). All the percentages are by weight. 1% (by weight; with respect to the monomers) of AIBN was added to each polymerization mixture.

| Column | AA% | MbA% | OctOH% | DMSO% | DMF% |
| --- | --- | --- | --- | --- | --- |
| A50 | 12.5 | 12.5 | 21.5 | 53.5 | - |
| A55 | 13.75 | 11.25 | 21.5 | 53.5 | - |
| A60 | 15 | 10 | 21.5 | 53.5 | - |
| A65 | 16.25 | 8.75 | 21.5 | 53.5 | - |
| 13DMF | 12.5 | 12.5 | 21.5 | 46.8 | 6.69 |
| 25DMF | 12.5 | 12.5 | 21.5 | 40.1 | 13.38 |
| 37DMF | 12.5 | 12.5 | 21.5 | 33.44 | 20.06 |
| 50DMF | 12.5 | 12.5 | 21.5 | 26.75 | 26.75 |

An attempt at scaling-up the column dimensions was made using glass-lined tubing of 0.8 mm ID (150 mm length). The tubing was silanized, vinylized and polymerized as described before. The A55 polymerization mixture was used. No further optimization was performed.

### PREPARATION OF PACKED CAPILLARY COLUMNS

The columns were packed by preparing a slurry of 0.1 g/mL 2.7 um AdvanceBio Glycan Mapping 120Å (Agilent,) particles using methanol as packing solvent. The slurry was inserted in an empty column (4.6 mm × 50 mm) with one end drilled out to be able to connect the capillary. The capillary (0.2 mm ID, 200 mm length; with a steel-based union and frits from VICI-Valco) was inserted 5 mm inside the packing column. A flow of 0.1 mL/min of the packing solvent was set. The column was kept under flow for 30 min after the packing was complete. The remaining slurry was collected and the packed column was removed and cut to a final length of 135 mm.

### INSTRUMENTS AND CONDITIONS

#### SEM experiments

Scanning electron microscopy (SEM) images of cross-sections of the monoliths were recorded on a FEI Verios 460 instrument (Thermo Fisher Scientific, Eindhoven, The Netherlands) equipped with an Everhart-Thornley detector (EDT) using a 2-kV electron beam. The samples were sputter-coated with a 15-nm gold layer on top of a 5-nm layer of palladium.

#### Permeability calculations

Permeability and hydrophilicity of the monolithic stationary phases were evaluated using a M-Class Acquity UPLC system (Waters, Milford, MA, USA), equipped with a binary solvent manager, a thermostated autosampler (1 μL sample loop) and a dual-wavelength tunable UV-vis detector with a 100 nL flow-cell. The permeability (*K_f_*) was determined by flushing MeOH through the acrylamide-based monolithic columns at various flow rates between 0.5 and 3 μL/min and using Darcy’s law

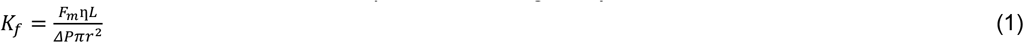

Where F_m_ is the flow rate of the solvent, η is the dynamic viscosity of the solvent, L is the length of the monolith, ΔP is the pressure drop across the monolithic column, and r is the radius of the capillary.

#### Hydrophilicity test

The hydrophilicity of the polymer monoliths was evaluated through the injection of a 0.1 mg/mL cytosine sample at different mobile-phases compositions. The sample was diluted in ACN and HP-water, matching the mobile phase composition. The separations were performed isocratically, at a flow rate 2 μL/min and at room temperature (22°C), with HP-water as mobile phase A and ACN as mobile phase B, both containing either 0.1% or 0.005% (v/v) of TFA.

#### Evaluation of monolith performance

Separations of intact proteins were performed on a UltiMate RSLCnano system (Thermo Fisher Scientific, Breda, The Netherlands) equipped with an autosampler (5 μL loop), thermostated column compartment with a ten-port, two-position valve and a loading-pump system (NCS-3500RS), and a UV-vis detector (VWD-3400RS). A trap-column (20 mm × 0.3 mm ID; C4 stationary phase, 5 μm particle size, 300 Å pore size; Thermo Fisher Scientific) was used to inject samples from water-based solutions, using a setup similar to that described by Gargano *et al*^34^.

Each reference protein was diluted in a solution of water : ACN = 95 : 5 with 0.1% TFA. The sample was loaded on the trap-column at 20 μL/min using the loading pump for 5 min with a mobile phase (A) of 2% ACN in water with 0.1% TFA. The valve was switched after 3 min from the beginning of the method.

Three washing cycles consisted of fast linear gradients from 20 to 80% and back to 20% of B (2% of HP-water in ACN) in 2 min were performed at the end of every separation, in order to avoid carryover. Further information about the method can be found in the figure captions.

The efficiency of the columns was assessed by calculating the peak capacity (*n*_c_) using the following equation (2):

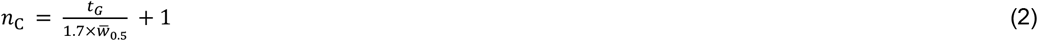

Where *t*_G_ is the gradient time and *w*_0.5_ is the average peak width at half-height.

The asymmetry of the peaks (A_*S*_) was calculated through the Chromeleon software (Thermo Fisher Scientific) according to the following equation (3):

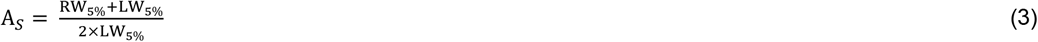

Where RW_5%_ and LW_5%_ are right and left peak width at 5% of the peak height, respectively.

The resolution for the separation of the five RnB proteoforms was estimated using the peak-valley ratio (P) equation (4) suggested by Christophe^35^:

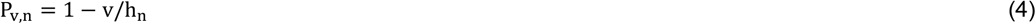

Where v is the signal intensity at the valley between two peaks, and h_n_ is the intensity of the peak taken into consideration (see explanation in Figure S6).

A separation of a protein mixture was performed also in the 0.8 μm ID monolithic column with the Waters M-class system described above. Further information about the chromatographic conditions and a protein separation (Figure S7) can be found in the SI.

#### HILIC-HRMS conditions

A Q-Exactive Plus Biopharma (Thermo Fisher Scientific, Bremen, DE) instrument was operated in positive-ion mode, using steel-based emitters, a nano-ESI source at a voltage of 2 kV, transfer capillary temperature of 300°C, and S-lens of RF 70.

The analyzed samples consisted of a 0.5 mg/mL solution of four proteins, *viz.* CC, CA, BSA and Tf, and a 0.5 mg/mL solution of RnB. The reference proteins were diluted in 10% ACN in HP-water. The volume injected was 0.5 μL for the protein mixture and 1 μL for the RnB sample.

Acquisition parameters for the protein mix where as follows. m/z range, 400 - 6000 m/z; resolution @200 m/z, 17,500; 10 microscans; max IT (Injection Time), 200 ms, AGC (Automatic Gain Control), 10^6^, in source CID (Collision-Induced Dissociation), 20 eV; HMR (High Mass Range) activated (trapping gas 1.2).

Acquisition parameters for RnB: m/z range 400 – 3000 m/z; resolution, 14000; 4 microscans; max IT 200 ms; AGC 10^6^, in source CID 20 eV, intact protein mode activated (trapping gas 0.2).

Total-ion current (TIC) and extracted-ion current (EIC) chromatograms and deconvoluted RnB masses were obtained using the Freestyle software (Thermo Fisher Scientific), while deconvoluted masses of the other proteins were obtained using UniDec^36^. Charge range, mass range and sample-mass interval were 1-30, 12,000-13,000 Da, 5 Da for CC, 1-50, 28,000-29,500 Da, 1.0 Da for CA, 1-50, 66,000-68,000, 1.0 Da For BSA, 1-80, 79,000-83,000 Da, 1.0 Da for Tf.

Extracted-ion current chromatograms of the RnB (140,000 Resolution) were obtained from 5 charge states (m/z 2129.3990, 1863.3501, 1656.4233, 1490.8819, 1242.5694) for RnB1, 5 charge states (m/z 2152.5490, 1883.6070, 1674.4294, 1507.1871, 1256.0739) for RnB2, 5 charge states (m/z 2175.6998, 1903.8632, 1692.4352, 1269.5785, 1523.2921) for RnB3, 5 charge states (m/z 2198.8515, 1924.1203, 1710.5526, 1539.5978, 1283.1666) for RnB4 and 5 charge state (m/z 2222.0021, 1944.5020, 1728.5585, 1555.7030, 1296.6706) for RnB5 with an extraction window of 10 ppm. A Gaussian-7 smoothing was applied to all the chromatograms. (MassIVE Repository link: ftp://massive.ucsd.edu/MSV000087977/)

## RESULTS AND DISCUSSION

### FABRICATION AND OPTIMIZATION OF THE MONOLITHIC STATIONARY PHASES

The composition of the polymerization mixture strongly affects the morphology and efficiency of organic monoliths. Based on the study of Xie *et al*^13^, we selected DMSO and OctOH as porogen system for our polymerization. While DMSO can dissolve the monomers (good solvent), it alone cannot provide sufficiently large pores. Therefore, OctOH is added as “poor solvent” to promote the development of flow-through pores.

In our experiments, we first used the ratio AA: MbA = 70: 30 described by Xie *et al.*^13^ and we polymerized the reaction mixture using DMSO/OctOH in 200-μm ID capillaries using thermal polymerization. The columns could be flushed with methanol. However, the polymeric material deteriorated when exposed to high amounts of water. Therefore, to obtain more rigid and water-resistant polymers, we used a higher percentage of crosslinker (AA: MbA = 50: 50), as described in Table 1. To guarantee an acceptable back pressure and a less-dense polymer structure, the overall amount of porogens employed was increased (from 70% in the original mixture to 75% w/w). Using these experimental conditions, we obtained polymer monoliths that could withstand pressures in excess of 40 MPa (400 bar; maximum pressure at which the columns were operated) under organic and aqueous conditions. Next we prepared a series of acrylamide-based monolithic columns to study the influence of the composition of the polymerization mixture, with particular focus on the effects of the AA-to-MbA ratio and the porogen system (using DMF as ternary solvent). The composition of the polymerization mixtures used is reported in Table 1.

### CHARACTERIZATION OF POLYACRYLAMIDE MONOLITHIC MATERIALS

To test the different acrylamide-based polymerization mixtures, we prepared capillary columns (triplicates for each polymerization mixture) and tested their permeability. Selected columns with different permeabilities were analyzed using scanning electron microscopy to verify the binding to the capillary wall and inspect the different globule features.

#### Permeability

The columns were tested before and after chromatographic separations and a change of permeability values, as well as a shift un the retention times was noticed for some of the columns (A60, A65, 13DMF and 50DMF). The permeability values of these columns are reported in SI (see Figure S1). It appears that a reduced amount of crosslinker (for A60 and A65) and early phase separation in case of ternary porogen mixtures (13DMF and 50DMF) can result in unstable monolithic structures that degrade rapidly.

According to Darcy’s law, the calculated permeability for the A50 columns, obtained from a 50/50 AA/MbA mixture was 4.4∙10^−14^ m^2^. Increasing the amount of crosslinker to 55% led to a small decrease in permeability (K_*f*_ = 5.4∙10^−14^). The lower permeability can be explained by an earlier phase separation during the polymerization, induced by a higher concentration of MbA. This leads to the formation of smaller pores. As far as the 25DMF columns are concerned, the introduction of a third non-polar porogen (DMF) in the polymerization mixture also appeared to induce an earlier phase separation compared to the binary porogen mixtures. However, the permeability of the resulting monolith decreased when adding 13 or 25% DMF and increased when increasing the DMF content to 37 or 50%, with the lowest permeability being observed for 25% DMF (K_*f*_ = 5.4∙10^−15^ m^2^). The latter is of the same order as that observed for 2.7-μm core-shell particles (K_*f*_ = 5.2∙10^−15^ m^2^) packed column.

#### Scanning Electron Microscopy

SEM images were used to confirm the attachment of the monolith to the capillary walls, the inspect the monolith homogeneity (Figure S2a-b), and to evaluate the influence of the different porogens, as well as the different concentrations of crosslinker used. We used this approach to analyze columns 25DMF, A50 and A55. All the monolithic stationary phases had a homogeneous micro-globular structure and a good attachment to the inner wall of the capillary. Figure 2 shows that the monolithic stationary phase in which 25% of the DMSO was substituted with DMF (column 25DMF, Figure 2A) features smaller globules than column A50 (Figure 2B) (approximately 0.5 μm compared to 1 μm). Similarly, Figure 2C (column A55) shows a small reduction of the globule size when increasing the percentage of crosslinker, which is in line with earlier observations of Viklund *et al*.^38^

**Figure 1.**
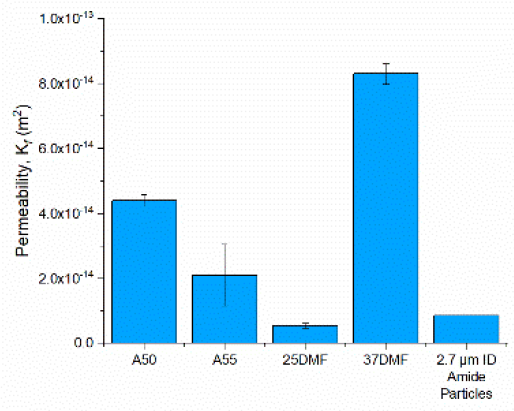
Permeability values of the acrylamide-based monolithic stationary phases.

**Figure 2.**
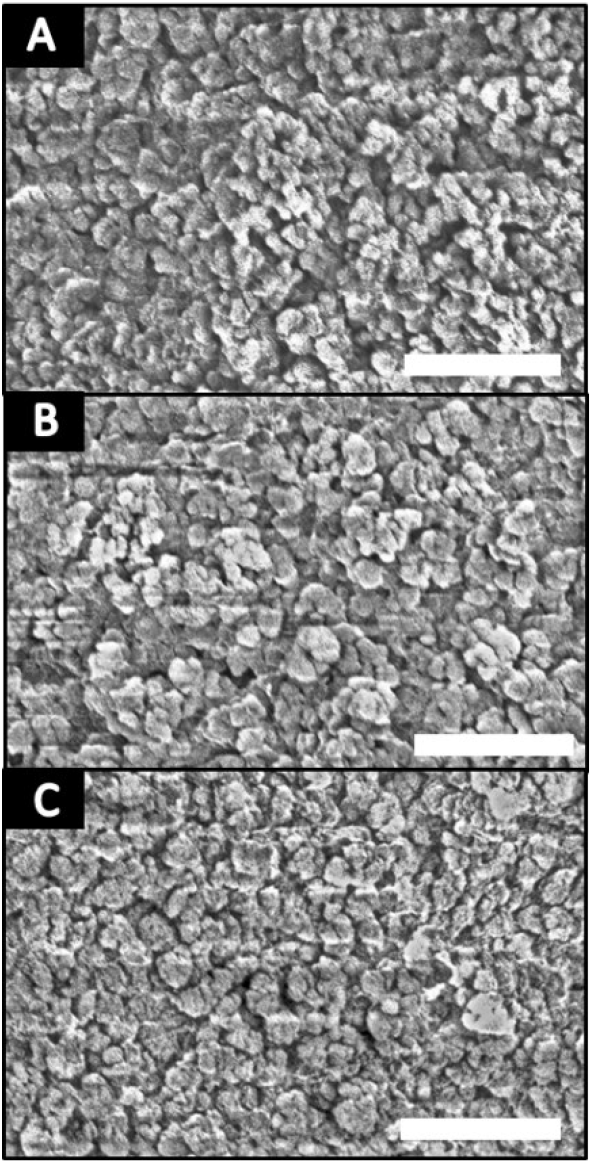
SEM images of cross-sections of three different acrylamide-based monolithic capillary columns. A) 25DMF, B) A50, C) A55. Scale bars 10 μm.

#### Hydrophilicity of the stationary phases: retention of nucleobases

An important characteristic of HILIC stationary phases is their hydrophilicity. The water intake of a polar stationary phase highly influences the retention mechanism and the chromatographic separation. To demonstrate that our monolithic columns show a typical HILIC behavior, we used A55 to perform isocratic separations of a polar compound (Cyt; Figure S3) at different water percentages. From this analysis it is clear that the stationary phase has mostly hydrophilic-interaction behavior (retention with ACN concentrations above 60%) and little or no RPLC retention (below 40% ACN), proving the hydrophilicity of the material.^39^

### CHROMATOGRAPHIC PERFORMANCE OF POLYACRYLAMIDE MONOLITHS

#### Polyacrylamide monolith for the separation of intact proteins

In order to find a good compromise between analysis time and efficiency, we tested a wide range of flow rates (from 1 μL/min to 5 μL/min) on an A55 capillary column, maintaining the gradient volume constant (70 μL, Table S2). The highest peak capacity (*n*_c_ = 106) was achieved at 1 μL/min in a 70-min gradient. However, to reduce the analysis time a flow rate of 2 μL/min was chosen and further used in our study.

To test the different column materials described in Table 1 we applied the same gradients (see experimental conditions) at 2 μL/min and 60°C. The columns were used to separate a mixture of reference proteins, *i.e.* Myo, Lys, CA and RNA. These proteins have different MW (13 to 30 kDa), pI (6.4 to 9.32) and low occurrence of proteoforms that could lead to peak broadening, due to partial resolution. In addition, we used RnB as a sample representative of glycoprotein separations. The results of this study are reported in Figure 3, S4 and S5.

**Figure 3.**
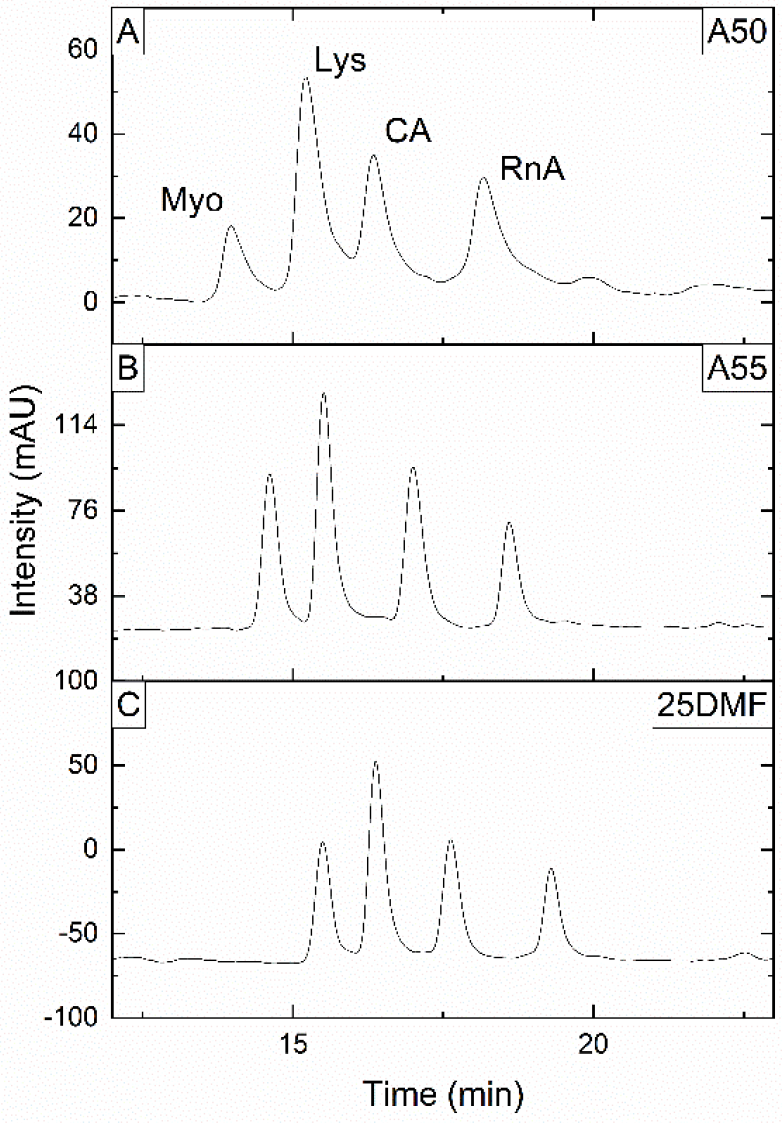
HILIC separation of intact proteins (A-C) on three capillary columns (A50, A55, 25DMF, top to bottom). The analysis was performed using a gradient from 90 to 85% (v/v) B in 1 min and then down to 55% B in 25 min at a flow rate of 2 μL/min. The analysis was carried at 60°C and the UV wavelength monitored was 214 nm. The chromatograms obtained with all the columns in table 1 are shown in Figure S4 of the SI.

The peak capacities for the protein mixture vary between 28 on the 13DMF column to 51 on the A55 column (Table S4). The columns present sufficient separation capacity for the RnB glycoforms. The five peaks are well-recognizable in the chromatograms obtained with most of the columns (Figure S5) and partially resolved (Table S5).

Figure 3 shows the separation of four proteins using the monolithic stationary phases that gave the best results, *i.e.* A50, A55 and 25 DMF. Good peak shape (with minor tailing; asymmetry between 1.13 and 1.47) and good peak capacities (between 36 and 51) were achieved on these columns. The lower permeability and higher retention times of the 25DMF column in comparison with the A55 columns (Table S3) suggest an increased surface area in the former column. Similar peak-valley-ratio values were found for the A55 and 25DMF columns (Table S5). All these three polymerization mixtures yielded good batch-to-batch repeatability (variation of retention time < 5%). Even though the A55 and 25DMF columns gave similar results, we chose A55 to continue our study, because of its better repeatability, higher permeability, and, consequently, the potential for the preparation of longer columns.

#### Comparison of protein separations on polyacrylamide monoliths vs. silica particles at different % of TFA

Next, we compared the performance of our polyacrylamide column A55 and a packed capillary column, to assess whether our newly developed material can be used with lower amounts of TFA. The packed column was prepared using 2.7-μm core-shell amide silica particles, as described in our previous work^25,28,31^. The two HILIC columns were compared using different concentrations of TFA (0.1 and 0.005% TFA) in the mobile phase (Figure 5). To evaluate the performance of the two columns we used a protein mixture that included CC, CA, BSA and Tf, and the glycoprotein RnB (Figure S8). These proteins cover a wide range of molecular weights (12 to 80 kDa) and isoelectric points (5.60 to 9.59).

**Figure 5.**
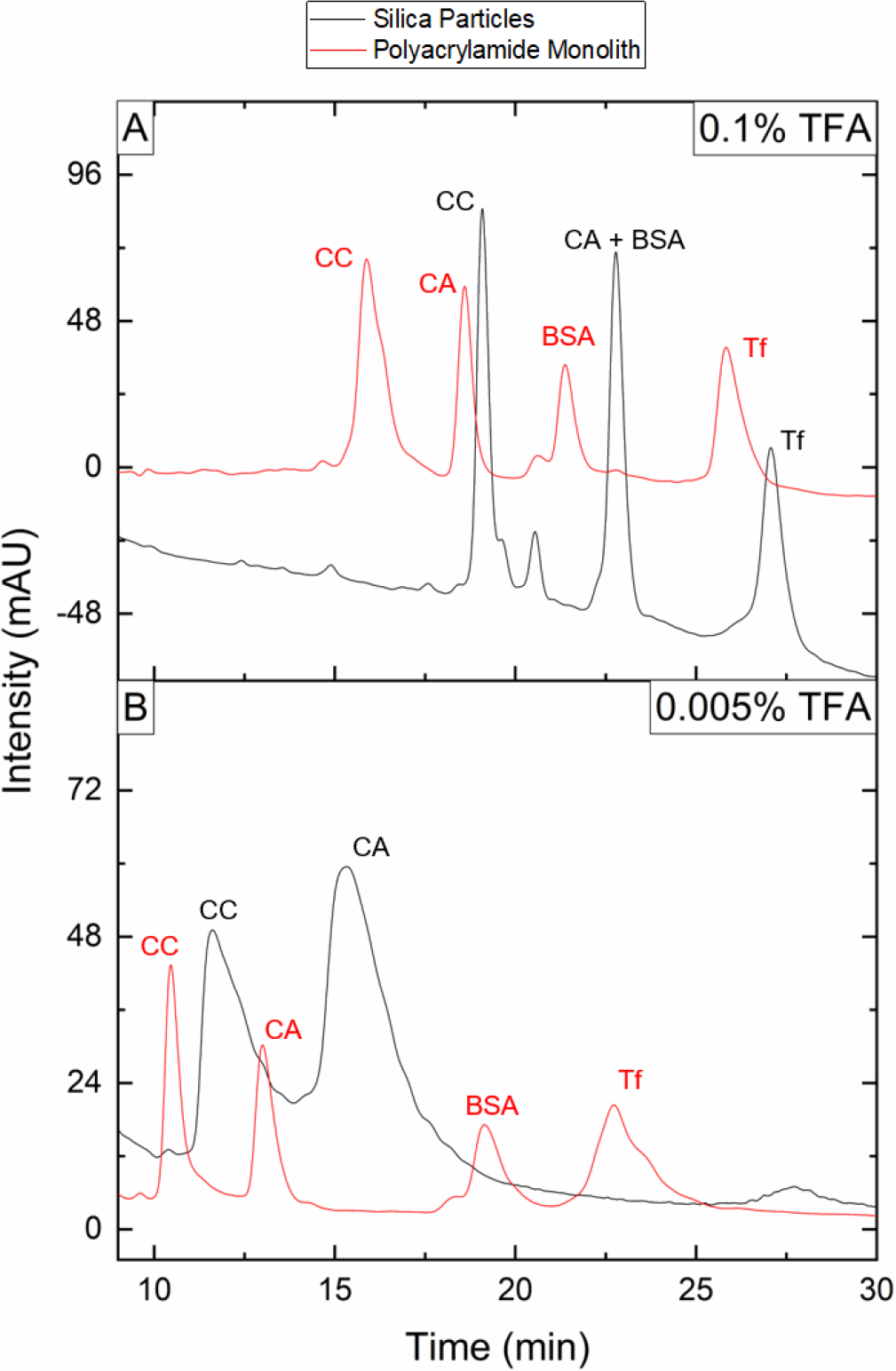
Comparison of HILIC separations of four intact proteins on the particle-packed column (black) and on the monolithic column (red). The analyses were performed using a gradient from 94 to 83% (v/v) of B in 1 min and then down to 65% B in 26 min with 0.1% (v/v) TFA (A) and with 0.005% (v/v) TFA (B) in the mobile phases. Protein elution order: CC, CA, BSA, Tf. All the analyses were carried at 45°C using a flow rate of 2 μL/min and the monitored UV wavelength was 214 nm.

We applied the same gradient (83-65% of B in 26 min) with a constant concentration of 0.1% (v/v) TFA for the separation of the protein mix on both columns. Interestingly, the peak capacity of the two columns was similar (32 vs 31), but the selectivity was different. On the (packed) silica-based column the protein mix was not fully separated and CA and BSA co-eluted, while on the polyacrylamide (monolithic) column the four peaks were baseline separated (Figure 5a and Table S6). The separation of RnB under gradient conditions (76-62% of B in 26 min) led to the partial separation of the five glycoforms on both columns with similar results in terms of peak-valley ratios (Table S7). Comparing the retention of the proteins between the two columns, the proteins eluted at a higher percentage of B (ca. 2% more ACN) from the monolithic column, possibly due to the lower surface area of this support (Table S6-7).

The same gradient used with 0.1% TFA was used for separations of the protein mixture with 0.005% TFA (Figure 5B). In the case of the polyacrylamide column, the retention decreased when moving from a high (0.1%) to a low concentration (0.005%) of TFA, with a minor reduction in the peak capacity (31 vs. 22). A possible explanation for the change in retention is the increased hydrophilic character of the water phase, due to the twenty-fold decrease in the concentration of the fluorinated ion-pairing reagent in the mobile phase. In the case of the packed silica-based column, one of the proteins (Tf) did not elute from the column (not even when increasing the water content up to 90%) and the separation significantly worsened (peak capacity of 12), with only two, significantly tailing peaks present in the chromatogram. We speculate that an increase in ion-exchange interactions, due to the decreased concentration of ion-pairing reagent in the mobile phase, may be responsible for this phenomenon. An in-depth analysis of the effects of different ion-pair agents and their concentrations is currently ongoing.

For the separation of RnB (Figure S8) we broadened the gradient in the case of the polyacrylamide column (83-63% B in 26 min), while a shallower gradient (75-68% B in 26 min) was used on the silica-based column. At a low concentration of ion-pairing reagent (0.005% TFA), RnB eluted at 75% and 73% B from the polyacrylamide and particle-packed columns, respectively. Lower percentages (71 and 69, respectively) were required when the concentration of TFA was 0.1% (v/v). Under optimal conditions for both columns, the monolithic column outperformed the packed silica-based column at 0.005% (v/v) TFA, as evidenced by the peak-valley ratio listed in Table S8.

#### Polyacrylamide columns for HILIC-HRMS at low TFA content

High-resolution mass spectrometry (HRMS) is an effective tool to characterize proteoforms of proteins, as these can be clearly identified based on mass shifts, with further confirmation by fragmentation experiments^8,40^. HILIC mobile phases are rich in ACN and this typically favours desolvation. However, in case of HILIC-MS of intact glycoproteins, the presence TFA (0.1% v/v, 13 mM), which is typically needed to ensure good chromatographic resolution of the glycoforms, causes signal suppression and protein-TFA adducts. The latter makes the mass spectra more complicated and more difficult to interpret. To reduce adducts formation high activation energies have to be used^28^. However, under these conditions some proteins may undergo fragmentation. For this reason, we have tested post-column solutions to decrease adduct formations during HILIC-MS in other research. ^34,41^ Reducing the content of TFA in the mobile phase should reduce adduct formation in HILIC-MS. To investigate this, we tested our polyacrylamide columns using mobile phases containing 0.1% TFA, 0.005%TFA and a combination of 0.1% FA and 0.005% TFA. We applied the same gradient (87-70% of B in 35 min) for the separations at lower concentration of TFA (0.005%), with and without FA.

Figure 6 shows TIC and EIC chromatograms recorded using the methods described in the captions at relatively low isCID (20 eV). The EIC chromatograms of the separation performed at higher TFA concentrations showed lower signal intensities and smaller peak areas (Table S8) in comparison with separations carried out with lower amouts of TFA. At high concentrations of TFA, the efficiency of the column was indeed higher than at a lower percentage of TFA (peak capacity, 42 vs. 15). The latter chromatogram showed broader peaks that result in a partial lost of resolution.

**Figure 6.**
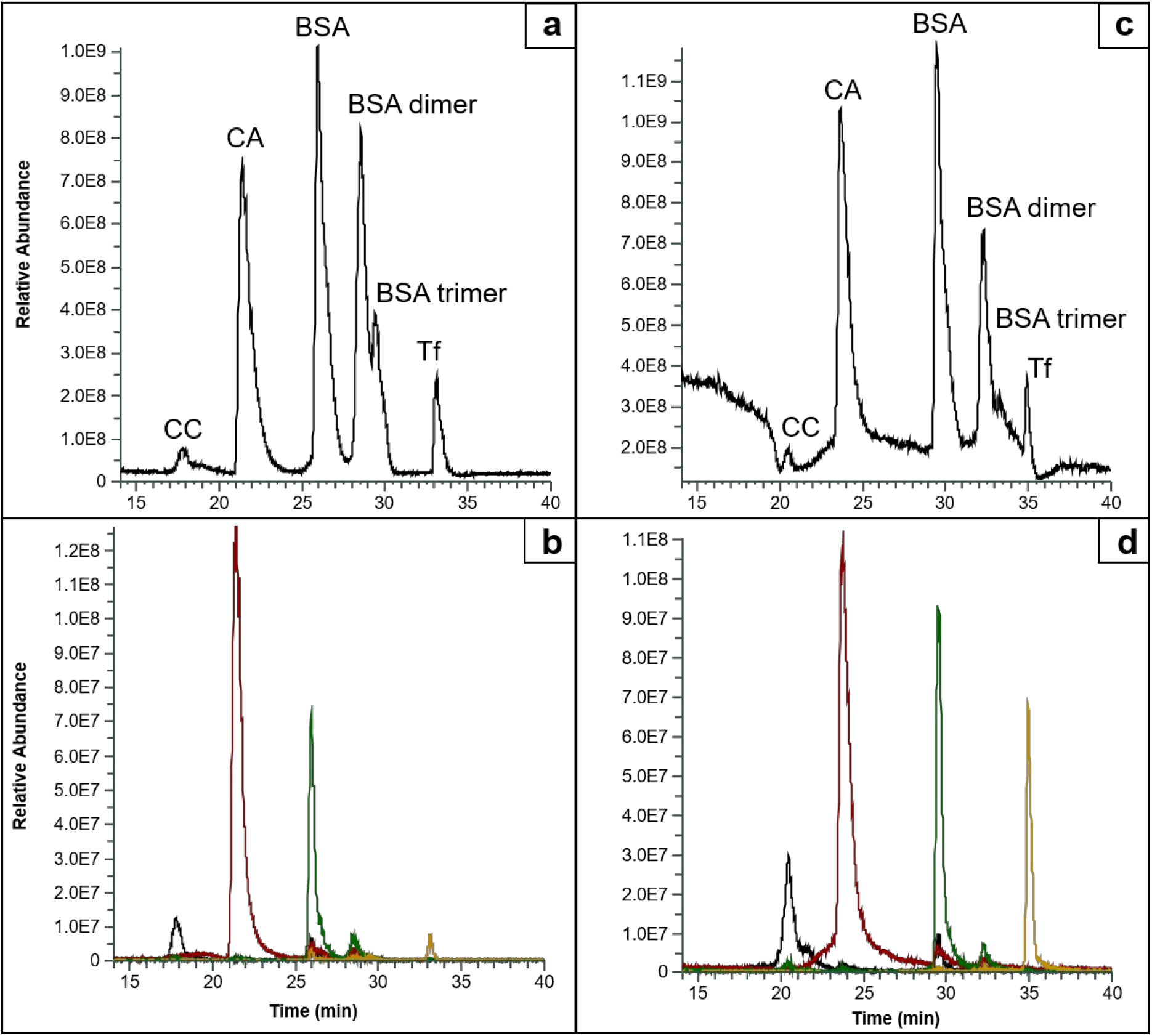
TIC (a,c) and EIC (b,d) chromatograms obtained using a HILIC-MS method with a, b) 0.1% (v/v) TFA, c, d) 0.005% (v/v) TFA + 0.1% (v/v) FA. A) gradient from 94 to 83% (v/v) of B in 2 min and then down to 67% in 35 min. B-D) gradient from 94 to 87% (v/v) of B in 2 min and then down to 70% in 35 min. All the analyses were carried at 45°C using a flow rate of 1.5 μL/min. The EIC chromatograms were obtained by summing the intensities of different charge state for CC (883.8, 951.7, 1030.9, 1124.4, 1236.8, 1374.1, 1545.7, 1766.2, 2060.5 m/z), CA (937.3, 968.4, 1001.9, 1037.6, 1076.6, 1117.3, 1161.9, 1210.4, 1262.8, 1320.3, 1383.0, 1452.3, 1528.5, 1613.4, 1708.2 m/z), BSA (1278.5, 1303.5, 1329.6, 1356.6, 1384.9, 1414.4, 1445.1, 1477.2, 1510.7, 1545.8, 1582.0, 1621.2, 1664.5, 1704.2 m/z) and Tf (2040.8, 2094.5, 2151.1, 2210.8, 2273.9, 2340.8, 2411.7, 2487.0, 2567.2, 2652.7, 2744.2 m/z). All the EIC are +/− 0.2 m/z

When adding FA to the mobile phase with low TFA content, the separation performances increased (peak capacity 40). Sharper peak shapes were obtained and the selectivity is restored. Importantly, FA did not influence the MS spectra and the signal intensity, since FA did not create adducts.

Figure 7 shows averaged spectra of the four proteins obtained from the HILIC-MS separations using different concentrations of TFA. The black traces indicate the protein spectra obtained using a mobile phases with 0.1% v/v TFA, while the red traces represent separations at a low concentration of TFA (0.005% (v/v) with 0.1% (v/v) FA added. The intensity of the red traces is higher than that of the black traces. This increase in sensitivity is most dramatic in the spectra of proteins with higher molecular weights (especially in Figure 7d). In fact, the signal intensity of the red trace for Tf is almost one order of magnitude higher than that of the black trace. The number of TFA-protein adducts is larger for the high-MW proteins than for the smaller proteins that have fewer interaction sites. Due to the formation of a lower number of adducts, a shift to higher charge states (lower m/z) is noticed when using 0.005% v/v TFA in the mobile phase. A lower amount of TFA reduces the charge distribution and shifts the most intense charge state. The corresponding deconvoluted specta reflect the trend in terms of adduct observed and are shown in Figure S9.

**Figure 7.**
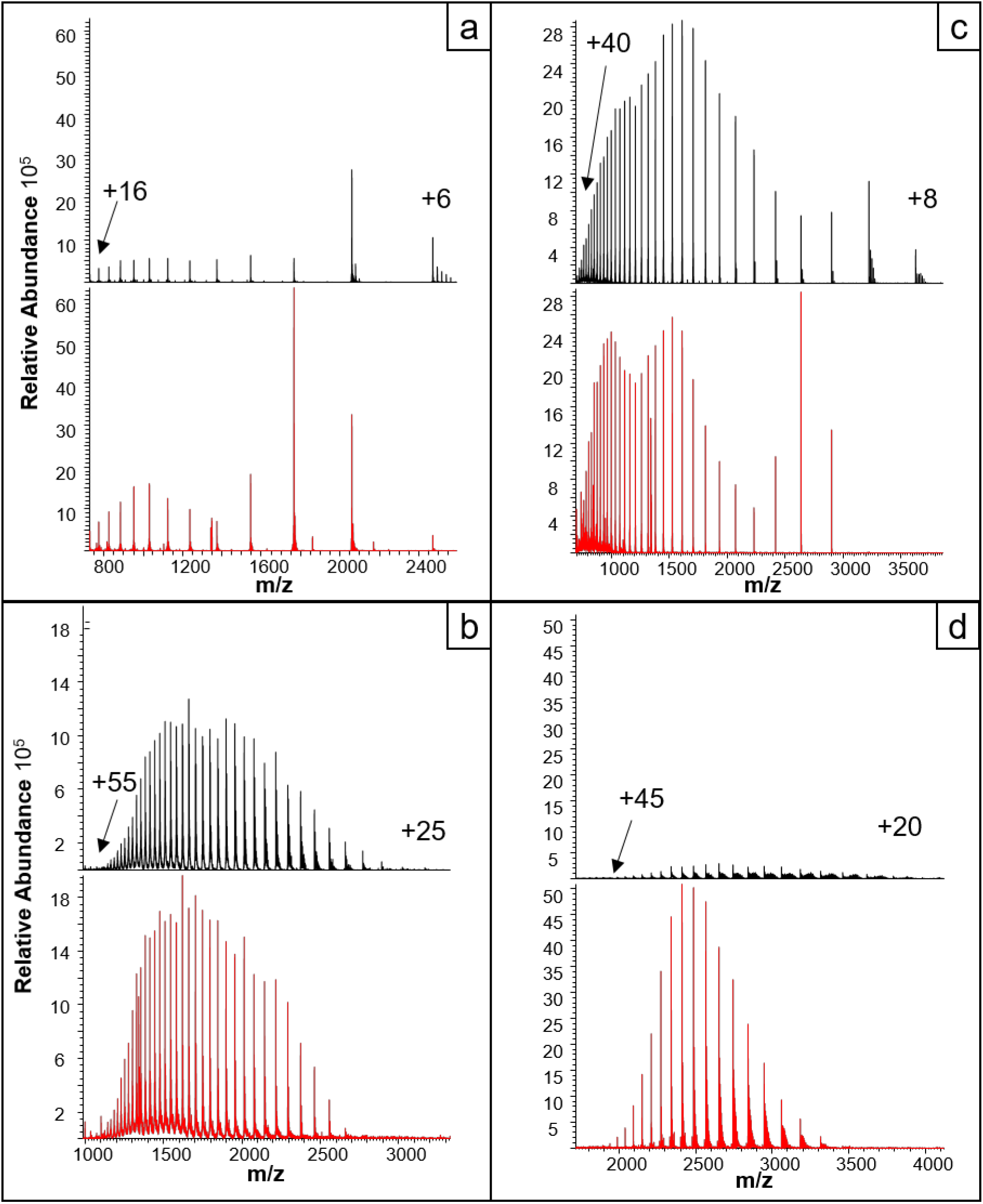
Average spectrum from the chromatographics peak of a) CC, b) CA, c) BSA, d) Tf obtained using the HILIC-MS method presented in Figure 6. 0.1% (v/v) TFA (black trace) and 0.005% (v/v) TFA + 0.1% (v/v) FA (red trace). The numbers on the mass spectra indicate the charge-state interval.

## CONCLUSIONS

Acrylamide-based monolithic stationary phases were successfully created within 200-μm ID capillaries using thermal polymerization. From SEM pictures, the micro-globular structure was seen to be homogenous and well bounded to the capillary wall. These highly hydrophilic monoliths were optimized for efficient HILIC separations of intact proteins. We selected the monolith with the polymerization mixture A55 as the best column, due to its high permeability and high efficiency (peak capacity 51).

At high concentrations of TFA (0.1% v/v), the A55 monolithic column showed greater selectivity, but a similar peak capacity as a packed (amide-modified silica) column. However, at lower concentrations of TFA (0.005%), the monolithic column showed no loss in selectivity and the peak capacity remained fairly high (31 to 22), whereas the packed column showed broader and more-asymmetrical peaks and a dramatic change in efficiency (peak capacity 32 to 12).

In coupling the separations with HRMS it was found that a reduction of the concentration of TFA in the mobile phase resulted in enhanced sensitivity, by one order of magnitude for the higher-MW proteins (e.g. Tf). The loss in chromatographic efficiency, incurred by reducing the TFA content, can be compensated by the addition of 0.1% (v/v) FA to the mobile phase on the monolithic column, which was found to raise the peak capacity from 15 back to 50. Moreover, from (deconvoluted) mass spectra it was clear that our polyacrylamide monolithic column showed a major advantage, as for high-MW proteins as a lower amount of TFA resulted in a lower number of TFA-protein adducts.

In future work we will try to exploit the polyacrylamide chemistry to create supports for stationary phases with a low hydrophobicity, for use in, for example, ion-exchange chromatography, affinity chromatography and size exclusion chromatography.

## SUPPORTING INFORMATION

The supporting information is available free of charge on the ACS Publication website at DOI:

## ACKNOWLEDGEMENTS

The STAMP project is funded under the Horizon 2020, Excellent-Science programme of the European Research Council (ERC), Project 694151. The sole responsibility of this publication lies with the authors. The European Union is not responsible for any use that may be made of the information contained therein.

